# PME-mediated pectin modifications promote haustoria initiation and xylem bridge development in the parasitic plant *Phtheirospermum japonicum*

**DOI:** 10.1101/2023.01.30.526340

**Authors:** Martina Leso, Anna Kokla, Ming Feng, Charles W. Melnyk

**Author notes:** **Corresponding author** Charles W. Melnyk, Department of Plant Biology, Linnean Center for Plant Biology, Swedish University of Agricultural Sciences, Almas allé 5, 756 51, Uppsala, Sweden.

## Abstract

- Parasitic plants produce cell wall modifying enzymes that are thought to be important for efficient host infection. Here, we investigated the role of pectin methylesterases (PMEs) and their inhibitors (PMEIs) during haustorium development in the facultative parasitic plant *Phtheirospermum japonicum* infecting *Arabidopsis thaliana*.
- We employed immunohistochemistry to characterise tissue-specific changes in pectin methylesterification during haustorium development. We found putative *PME* and *PMEI* genes in *P. japonicum* and used genetic and transcriptomic approaches to identify those involved in haustorium development.
- Our results show tissue-specific changes in pectin methylesterification during haustorium development. De-methylesterified pectin correlated with haustorial intrusive cells whereas highly methylated pectin correlated with vascular tissues. We also found that inhibition of PME activity delayed haustoria development and xylem connectivity. Several *PjPME* and *PjPMEI* genes increased expression specifically during haustorium development but such increases did not occur when haustorium initiation or xylem connections were blocked by chemical treatment.
- This study describes the importance of pectin modifications in parasitic plants during host infection. Our results suggest a dynamic regulation of PMEs and PMEIs contributes to haustoria initiation and to the establishment of xylem connections between parasite and host.

## Introduction

Parasitic plants, which constitute around 1% of angiosperm species (Westwood et al. 2010), are devastating pests that cause major agricultural losses each year (Rodenburg et al. 2016). Parasitism has evolved independently at least 11 times, and despite these diverse origins, all parasitic plants form an invasive structure, the haustorium, which penetrates the host and allows the uptake of nutrients, hormones and signalling molecules (Birschwilks et al. 2006; Spallek et al. 2017; Shahid et al. 2018; Liu et al. 2020). The development of the haustorium starts with the perception of a suitable host through haustorium inducing factors (HIFs). Treatment with 2,6-Dimethoxybenzoquinone (DMBQ), the first discovered HIF, is sufficient to induce the formation of pre-haustoria even in the absence of a host (Chang et al. 1986). Other HIFs include hormones like cytokinin and lignin-related compounds (Goyet et al. 2017; Cui et al. 2018; Aoki et al. 2022). In the facultative root parasite *Phtheirospermum japonicum* the perception of HIFs is mediated by leucine-rich-repeat receptor-like kinases (Laohavisit et al. 2020) and increases auxin polar transport and auxin biosynthesis. The auxin signalling peak promotes cell expansion and division, leading to the formation of a swelling called the pre-haustorium (Ishida et al. 2016; Wakatake et al. 2020). Penetration of the host by the pre-haustorium is thought to depend on haustorium-secreted cell-wall-modifying enzymes such as expansins and peroxidases that loosen the host cell walls (Losner-Goshen et al. 1998; Veronesi et al. 2007; Honaas et al. 2013; Olsen et al. 2016). The invasion of the host tissues is then mediated by the intrusive cells, which differentiate from epidermal cells at the parasite-host interface and drive haustorial growth towards the host vasculature (Heide-Jørgensen and Kuijt 1995; Hood et al. 1998). Finally, a vascular connection develops between the parasite and the host. All parasitic plants form a xylem connection, which begins its differentiation from the cambium-like tissue at the centre of the haustorium (Wakatake et al. 2018). A mass of xylem tissue then develops close to the parasite vasculature (plate xylem) before strands of xylem (xylem bridges) differentiate to connect the xylem of the parasite to the host.

Despite recent advances in our understanding of haustorium development, the mechanisms regulating haustoria initiation and host invasion remain largely unknown, but likely rely in part on cell wall modifications. In plants, lateral organ development relies on the fine tuning of cell wall modifications which are required for cell expansion, division and differentiation. These processes all require the modification of cell-to-cell adhesion. The main mediator of cell adhesion in plants is pectin, a jelly-like matrix composed of homogalacturonan, rhamnogalacturonan I and rhamnogalacturonan II (Pelloux et al. 2007; Daher and Braybrook 2015). Homogalacturonan is secreted to the cell wall in a highly methylesterified state and is then modified in the cell wall by different families of pectin modifying enzymes including pectin methylesterases (PMEs), PME inhibitors (PMEIs), polygalacturonases (PGs) and pectate lyases (PLs). Highly methylesterified pectin forms a tight matrix with less elastic properties. During developmental processes such as tissue expansion and lateral organ emergence, homogalacturonans undergo de-methylesterification by PMEs to become looser (Pelloux et al. 2007; Daher and Braybrook 2015). In addition to the fundamental roles of PMEs and PMEIs in plant development, many plant pathogens hijack plant pectin modification mechanisms to allow tissue intrusion (Hewezi et al. 2008; Raiola et al. 2011). For example, the cyst nematode *Heterodera schachtii* secretes into *Arabidopsis thaliana* a cellulose binding protein (CBP) that activates PME3, facilitating the entry of the nematode (Hewezi et al. 2008). Mutating *pme3* makes *A. thaliana* less susceptible to nematode infection (Hewezi et al. 2008). Furthermore, some nematodes and fungi can directly secrete PMEs that mimic plant PMEs and facilitate host tissues invasion (Valette-Collet et al. 2003; Vicente et al. 2019). Parasitic plants are thought to secrete cell-wall modifying enzymes (CWMEs) to allow both the growth of the haustorium as well as to loosen host tissues to allow for invasion. The secretion of CWMEs has been partially investigated in some parasitic plant species. Different species from the *Orobanche* genus secrete PMEs, PGs, PLs and peroxidases close to the site of infection to modify the host’s cell wall (Ben-Hod et al. 1993; Losner-Goshen et al. 1998; Veronesi et al. 2007), while the root parasite *Thesium chinense* upregulates *PME* transcription during haustorium development (Ichihashi et al. 2018).

Even though cell wall modifications are suggested to be crucial for haustorium development, this aspect remains largely unexplored. Here, we demonstrate that PME activity is important for haustoria development in the parasitic plant *P. japonicum*. We show that the degree of pectin methylesterification is regulated in a tissue-specific manner, and that chemical or genetic inhibition of the parasite’s PME activity impairs haustorium formation and xylem connectivity. Finally, we show that *PjPMEs* are expressed in intrusive cells while *PjPMEI* are mostly expressed in other tissues, suggesting a specialised function in specific stages of haustorium formation.

## Materials and Methods

### Plant material and growth conditions

*P. japonicum* and *A. thaliana* seeds were surface sterilised by washing with 70% ethanol for 20 minutes, followed by 95% ethanol for 5 minutes, and sown on 1⁄2 MS medium with 1% sucrose and 0.8% bactoagar. After stratification at 4°C in darkness for 1 or 2 days for *P. japonicum* and *A. thaliana* respectively, the plates were moved to a growth cabinet at 25°C in long day conditions (16h light/8h darkness), 100 μmol m^−2^ s^−1^ light. The *A. thaliana* Col-0 accession was used unless otherwise stated. The *AtPMEI5OE* line has been previously published (Wolf et al. 2012). The BR-signalling mutants *bes1-2* and *bes1D* have been previously published (Yin et al. 2002; Lachowiec et al. 2013).

### PjPMEs and PjPMEIs identification and phylogenetic analyses

*A. thaliana* PME (Louvet et al. 2006) and PMEI (Wang et al. 2013) sequences were downloaded from the Phytozome database (Goodstein et al. 2012). *P. japonicum* putative PMEs and PMEIs were identified by searching the HMM profiles (PF01095 and PF04043 respectively) on a *P. japonicum* proteome obtained from the published genome (Cui et al. 2020) using the HMMER3 software (Finn et al. 2011). The putative PjPMEs were the aligned using Clustal W in MEGAX (Tamura, Stecher, and Kumar 2021), and the sequences lacking more than one of the five conserved catalytic amino acids (Johansson et al. 2002; Markovic 2004) were removed from downstream analyses. *P. japonicum* and *A. thaliana* PME and PMEI sequences were aligned using ClustalW. Maximum-Likelihood phylogenetic trees were built using MEGAX with 100 bootstraps.

### *In vitro* infection assays with *Phtheirospermum japonicum*

Infection assays were performed according to Kokla et al. 2022. Briefly, five days after germination *P. japonicum* seedlings were moved from nutrient medium to nutrient-free medium (water agar) for starvation. After three days, a six-day old *A. thaliana* seedling was aligned root-to-root to each *P. japonicum* seedling to allow infection. 50 uM or 100 uM EGCG, 100 nM or 200 nM epiBL, 5 uM NPA, 0.05 mM Coomassie Brilliant Blue and 0 to 500 nM NAA were applied directly in the nutrient-free medium and left until the end of the infection. For measuring the plate xylem area and the number of xylem bridges, 7 days post infection (dpi) haustoria were stained with Safranin-O following Spallek et al. 2017. Pictures were taken using an Axioscope A1 microscope and analysed in Fiji (Schindelin et al. 2012; Rueden et al. 2017).

### Immunohistochemical staining of pectin residues

*P. japonicum* infecting *A. thaliana* was harvested at 0, 24, 48, 72 and 120 hpi for infections on water, and at 0 and 72 hpi for infections on DMSO, 100 nM epiBL, 0.05 mM Coomassie and 5 uM NPA treatments. The seedlings were fixed in a 1% glutaraldehyde, 4% formaldehyde, 0.05 M NaPI aqueous solution by vacuuming twice for 20 minutes, followed by overnight incubation at 4°C. The samples were then dehydrated with an ethanol gradient (30 minutes in each of 10%, 30%, 50%, 70%, 96%, 100%, 100% ethanol) and incubated overnight in a 1:1 solution of 100% ethanol:Historesin solution (Leica). The solution was exchanged with Historesin and the samples incubated again overnight at 4°C. The seedlings were then oriented in molds following (Scheres et al. 1994), aligning the haustoria of different seedlings. The shoot was removed, and a 14:1 solution of Historesin and hardener was added to form a hard resin sheet. The haustoria were cross-sectioned at 8 um thickness using a Microm HM355 S microtome. The sections were rehydrated in PBS, incubated in 0.05 M glycine in PBS for 20 minutes, and blocked in 2% BSA in PBS (blocking buffer) for 30 minutes. Three consecutive slides were stained in 1:20 dilutions of LM19, LM20 in PBS or just PBS for the negative control, and incubated for 2 h. After rinsing with blocking buffer three times, the sections were incubated for 1 h in a 1:100 dilution of Goat anti-Rat IgG Alexa Fluor 647 secondary antibody. After rinsing three times with PBS, the sections were mounted in PBS and immediately imaged on a Zeiss LSM-780 confocal microscope. Images were processed in Fiji using the 16-colors LUT.

### Ruthenium red staining

Ruthenium red staining was performed by dipping infecting roots at 0, 24, 48, 72 and 120 hpi in 0.05% ruthenium red in deionised water for 5 minutes, followed by rinsing 2 times with deionised water and mounting on 20% glycerol. Pictures were taken using an Axioscope A1 microscope and analysed in Fiji (Schindelin et al. 2012; Rueden et al. 2017).

### RNA-seq datasets and genes accession numbers

The RNA-seq dataset used for gene expression analyses of the infection time course in *P. japonicum* and *A. thaliana* is presented in Kokla et al. 2022. The dataset used for gene expression analyses in intrusive and non-intrusive haustorial cells is presented in Ogawa et al. 2021. Genes IDs and accession numbers for the *P. japonicum* genes mentioned in the text are available in Supplementary Table 2.

### Gene expression analyses

40 five days old *P. japonicum* seedlings per sample were transferred to the starvation medium with 100 uM EGCG, 100 nM epiBL, 5 uM NPA, 0.05 mM Coomassie, 1 uM NAA for three days or control DMSO. After infection with *A. thaliana*, 2 mm of root around the haustorium was collected at 0 or 72 hours post infection. RNA was extracted using the ROTI®Prep RNA MINI kit (Carl Roth®, 8485) following the manufacturer’s instructions. cDNA was synthesised with the Maxima First Strand cDNA Synthesis Kit for RT-qPCR (ThermoFisher, K1642) following the manufacturer’s instructions. qRT-PCR was performed using the Maxima SYBR Green/ROX qPCR Master Mix 2x (Thermo Scientific, K022). *PjPP2A* was used as normalisation control (Serivichyaswat et al. 2022). For each experiment, three biological replicates and at least two technical replicates were used. The relative gene expression was calculated using the Pfaffl method. The primers used are available in Supplementary Table 3.

### Cloning of *PjPMEs* and *PjPMEIs* and plasmid construction

All cloning was based on the Greengate cloning method following the standard protocols (Lampropoulos et al. 2013). For the overexpression constructs, the CDS of *PjPME6, PjPME51*, *PjPMEI9, PjPMEI6* and *PjPMEI10* were amplified using the CloneAmp™ HiFi PCR Premix (TakaraBio) from the cDNA of *P. japonicum* and inserted into the entry vector pGGC000. The ligated plasmids were amplified in chemically competent *E. coli* DH5α and confirmed by Sanger sequencing. The final binary vector assembly was performed using pGGA-pMAS, pGGB0003, pGGC-CDS, pGGD002, pGGE-terMAS, pGGF-DsRed and pGGZ001. pGGA-pMAS, pGGE-terMAS and pGGF-DsRed were previously published (Kokla et al. 2022). For the reporter constructs, a sequence of ~3 kb upstream the starting codon was cloned as the promoter of the genes of interest (GOI). *PjPME6* (3087 bp), *PjPME51* (2876 bp), *PjPME22* (3044 bp), *PjPMEI9* (3021 bp), *PjPMEI10* (3071 bp) and *PjPMEI6* (2988 bp) promoters were amplified using the CloneAmp™ HiFi PCR Premix (TakaraBio) from the gDNA of *P. japonicum* and inserted into the entry vector pGGA000. A 2xVenus-NLS sequence was cloned from a previously published GoldenGate vector backbone (Cui et al. 2016) and inserted in the pGGC000 entry vector to create pGGC-2xVenus-NLS. The ligated plasmids were amplified in chemically competent *E. coli* DH5α and confirmed by Sanger sequencing. The final binary vector assembly was performed using pGGA-pGOI, pGGB0003, pGGC-2xVenus-NLS, pGGD002, pGGE001, pGGF-DsRed and pGGZ001. The final overexpression and reporter plasmids were co-transformed in electrocompetent *Agrobacterium rhizogenes* AR1193 with the pSoup plasmid, and the bacteria cultured in LB broth with 50 ug/ml spectinomycin and 50 ug/ml rifampicin.

### *P. japonicum* hairy root transformation

*P. japonicum* transformation was performed according to Ishida et al. 2011. Seven-day-old *P. japonicum* seedlings were sonicated for 10 seconds and vacuum-infiltrated for 5 minutes in a solution of AR1193 carrying the construct of interest. The seedlings were then moved to solid B5 medium supplemented with 1% sucrose and 450 uM acetosyringone and kept at 22°C in the dark for 2 days. Seedlings were then moved to B5 medium containing 300 ug/ml cefotaxime and grown at 25°C in long days conditions until formation of hairy roots. Transgenic hairy roots were identified through red fluorescence using a Leica M205 FA stereo microscope and placed on starvation medium for 4 days before addition of *A. thaliana*. Non-fluorescent hairy roots from the same transformation experiment were used as a control for each construct. Counting of haustoria and safranin-O staining were performed at 7 dpi for overexpression constructs. Imaging of transcriptional reporters was performed on 4 dpi haustoria using a Zeiss LSM780 confocal microscope with 514 nm excitation, 2.8% laser power and 519-550 nm detection.

### Statistics

All experiments were replicated at least three times unless otherwise stated. For infection assays each biological replicate consisted on the average of results from at least 15 plants, and two-tailed student’s t-tests on means were used for single comparisons. For infection of transformed hairy roots overexpressing *PjPMEs* or *PjPMEIs*, the data from the biological replicates were pooled and divided in categories of 0, 1 or >= 2 haustoria per hairy root. A Fisher exact test was then used to calculate significance. For qPCR data, two-tailed student’s t-tests on biological replicates were used for single comparisons of treatment VS control.

## Results

### *PjPMEs* and *PjPMEIs* are differentially expressed during haustorium development

To study the role of pectin modifications in parasitism of *A. thaliana* by *P. japonicum*, we focused on pectin methylesterase (PME) and PME-inhibitor (PMEI) enzyme families. We performed a Hidden-Markov-Model search on the *P. japonicum* (Pj) proteome (Cui et al. 2020), and identified 73 putative PjPMEs and 62 PjPMEIs (Supplementary Table1). We further filtered PjPMEs based on the presence of at least three of the five conserved catalytic amino acids (Johansson et al. 2002; Markovic 2004) and retained 60 PjPMEs for downstream analyses (Supplementary Table1). We aligned PjPMEs and PjPMEIs with *A. thaliana* PMEs and PMEIs respectively, and built two Maximum-Likelihood phylogenetic trees. The trees obtained show co-clustering of the sequences from *P. japonicum* with the ones from *A. thaliana*, suggesting conservation in PMEs and PMEIs sequences between parasite and host (Supplementary Fig. 1a-b). We then looked at the expression of *PjPMEs* and *PjPMEIs* in two different *P. japonicum* transcriptomic datasets. The first dataset sampled tissues at the site of haustorium development during a time-course infection of *A. thaliana* (Fig. 1a, Kokla et al. 2022), while the second dataset sampled intrusive cells (ICs) and non-ICs in mature haustoria infecting rice (*Oryza sativa*) (Fig. 1b, Ogawa et al. 2021). RNA levels of several *PjPME* and *PjPMEI* genes highly increased during haustorium formation, particularly at 48 and 72 hours post infection (hpi), while very few genes showed decreased RNA levels (Fig. 1c,d). Most of the *PjPME* genes that increased expression in the time-course dataset showed higher expression in ICs compared to non-ICs, while *PjPMEIs* were more highly expressed in non-ICs compare to ICs (Fig. 1c,d). In the *A. thaliana* host dataset, most *AtPMEs* and *AtPMEIs* did not show clear increased or decreased expression during haustorium development, with *AT1G23200* (*PME*) and *AT2G01610* (*PMEI*) showing the most consistent pattern of increased expression over multiple time points (Supplementary Fig. 1c,d).

**Figure 1:**
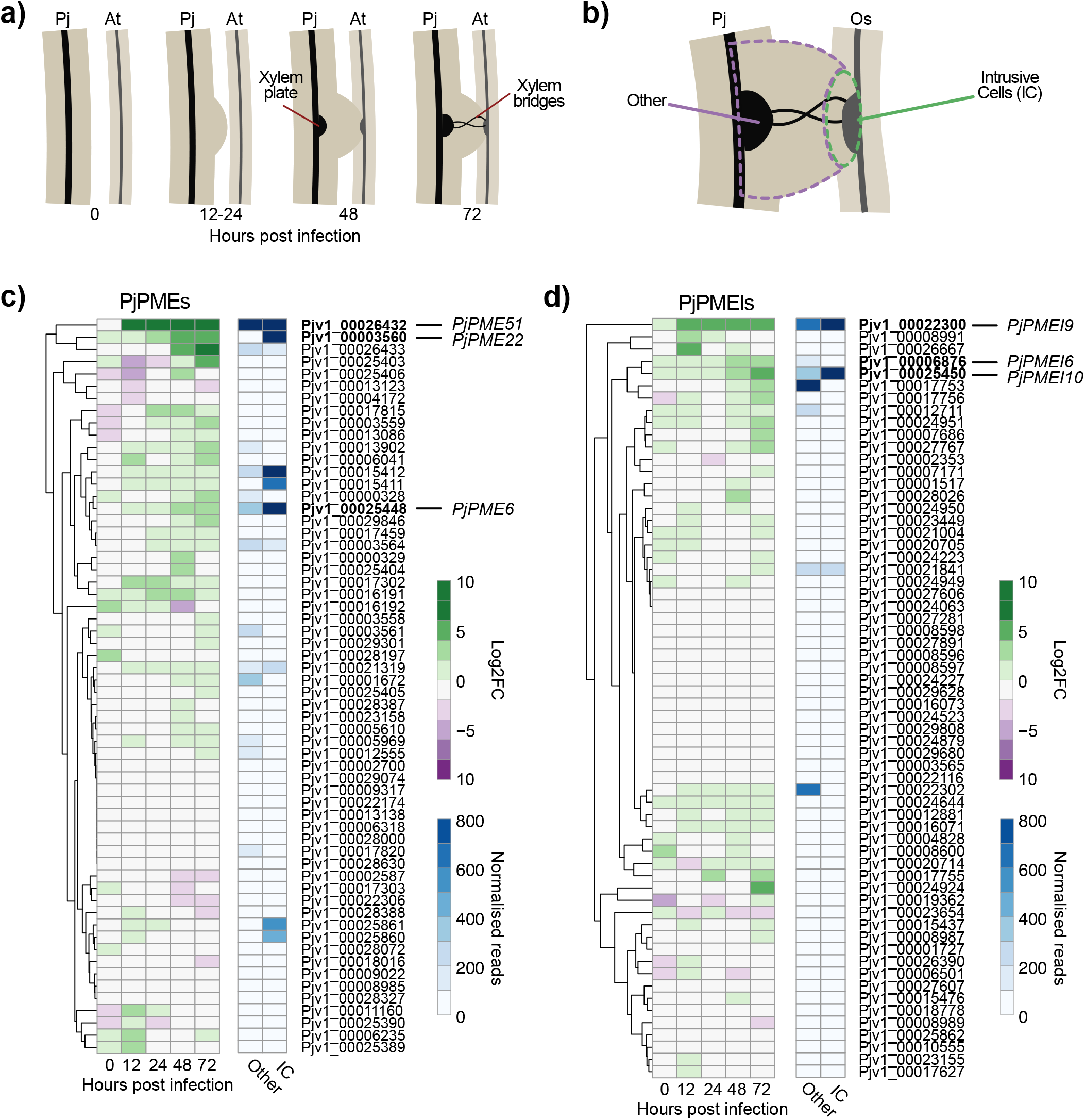
*PjPMEs* and *PjPMEIs* are differentially expressed during haustorium development. a) Illustration of *Phtheirospermum japonicum* haustorium development during infection of *Arabidopsis thaliana* corresponding to the time points selected for the time course RNA-seq (Kokla et al. 2022) b) Illustration of the intrusive cells (IC) or rest of the haustorial tissues (other) corresponding to the intrusive cells RNA-seq (Ogawa et al. 2021). c) Heatmaps of the expression of candidate *P. japonicum PMEs* (*PjPMEs*): log2 fold change between *P. japonicum* infecting and not infecting over five time points during infection, and normalised reads in intrusive cells and other haustorial tissue. d) Heatmaps of the expression of candidate *P. japonicum PMEIs* (*PjPMEIs*): log2 fold change between *P. japonicum* infecting and not infecting over five time points during infection, and normalised reads in intrusive cells and other haustorial tissue. Pj = *Phtheirospermum japonicum*, At = *Arabidopsis thaliana*, Os = *Oryza sativa*

### PME activity increases during haustorium development and is higher in intrusive cells

Since we observed differential expression of *PjPMEs* and *PjPMEIs* during haustorium development, we tested whether pectin methylesterification levels could also be affected. We first performed ruthenium red staining on whole roots during an infection time course (Fig. 2a). Ruthenium red binds with higher affinity to de-methylesterified homogalacturonan, and therefore a higher staining often correlates with higher PME activity (Downie et al. 1998). We observed an increase in ruthenium red staining by 24 hpi at the interface between *P. japonicum* and *A. thaliana*. The staining intensity increased further in later stages corresponding to host invasion (48 hpi) and xylem differentiation (72 and 120 hpi) (Fig. 2a,b, Supplementary Fig. 2a), but was not observed when inducing haustoria using DMBQ (Fig. 2b, Supplementary Fig. 2b), which does not induce xylem bridge formation. We also used LM19 and LM20 antibodies, specific for de-methylesterified and highly methylesterified pectin respectively, to measure pectin modifications from 0 to 120 hpi (Fig. 2c, Supplementary Fig. 2c). At early haustorium development stages (24 hpi), the epidermis showed highly demethylesterified pectin via LM19, while inner tissues showed highly methylesterified pectin via LM20. At 48 hpi, LM19 staining was strong in intrusive cells and in the developing cambium-like tissues, while staining with LM20 was reduced. In completely developed haustoria (120 hpi), we saw an absence of LM19 staining in the xylem bridge, with the surrounding tissues more strongly stained by LM20 (Fig. 2c,d). These results showed that pectin methylesterification is highly plastic and is modified dynamically during haustorium development in a tissue-specific manner.

**Figure 2:**
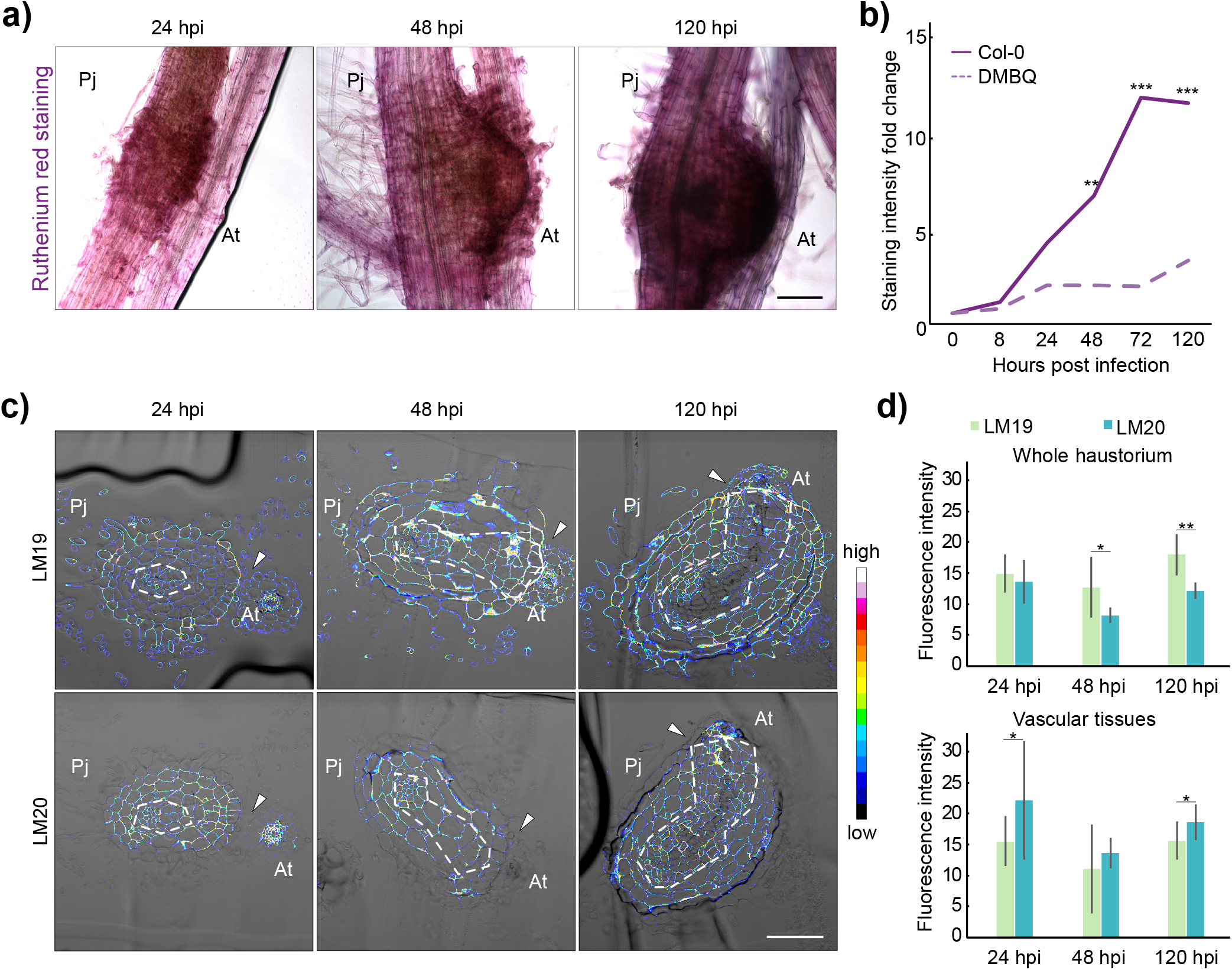
PME activity increases during haustorium development and is higher in intrusive cells. a) Ruthenium red staining of the developing haustoria at 24, 48 and 120 hours post infection (hpi). b) Quantification of staining intensity in infecting haustoria (solid line) or pre-haustoria formed on DMBQ (dashed line) during a time course from 0 to 120 hpi. Staining intensity normalised to 0 hpi. Asterisks indicate significant increase in staining compared to 0 hpi (Student’s t-test). c) Fluorescence images of antibody staining with LM19 (unmethylated homogalacturonan) and LM20 (highly methylated homogalacturonan) on haustoria cross sections at 24, 48 and 120 hpi. Vascular tissues are circled in white dashed lines. d) Fluorescence quantification in whole *P. japonicum* sections and vascular tissues. Stars indicate significant difference between LM19 and LM20 staining intensity per time point (Student’s t-test). Scale bars 100 um; Pj = *Phtheirospermum japonicum*, At = *Arabidopsis thaliana*; arrowheads denote the parasite-host interface. * for p<0.05, ** for p<0.01, *** for p<0.001.

### Differentially expressed *PjPMEs* and *PjPMEIs* express in intrusive cells and cambium-like tissue

Since the pectin degree of methylesterification (DM) in the developing haustorium was different in specific tissues, we investigated candidate *PjPMEs* and *PjPMEIs* to understand if their expression pattern correlated with the pattern of pectin DM. We chose three *PjPME* genes with increased expression in the time course dataset and renamed them based on the closest *A. thaliana* homolog (Fig. 1c, 3a). All three genes were also upregulated during *P. japonicum*-*O.sativa* infection. *PjPME6* (*Pjv1_00025448*) and *PjPME22* (*Pjv1_00003560*) expression levels increased in ICs compared to other tissues, while *PjPME51* (*Pjv1_00026432*) was highly expressed in both IC and non-IC tissues (Fig. 3b). We made and transformed transcriptional reporters in *P. japonicum* hairy roots and found *PjPME6* and *PjPME51* reporters showed signal mainly in ICs, while the *PjPME22* reporter was mostly expressed in *P. japonicum* vasculature (Fig. 3c). We then selected three *PjPMEI* genes upregulated during haustorium development (Fig. 1d, 3d). In the *P. japonicum*-*O.sativa* dataset, *PjPMEI9* and *PjPMEI10* showed increased expression in ICs, while *PjPMEI6* was more expressed in non-IC tissues (Fig. 3e). Our transcriptional reporters for these genes showed expression in ICs and cambium-like tissues for *PjPMEI9*, and plate xylem and ICs for *PjPMEI10* (Fig. 3f). The *PjPMEI6* reporter showed no fluorescence in 4 dpi haustoria (Fig. 3f). All the *PjPMEs* and *PjPMEIs* we investigated were not expressed in the primary root tip of the hairy roots, and showed little or no fluorescence at the lateral root emergence sites of the hairy roots, suggesting that upregulation of these genes might be specific to haustorium development (Supplementary Fig. 3).

**Figure 3:**
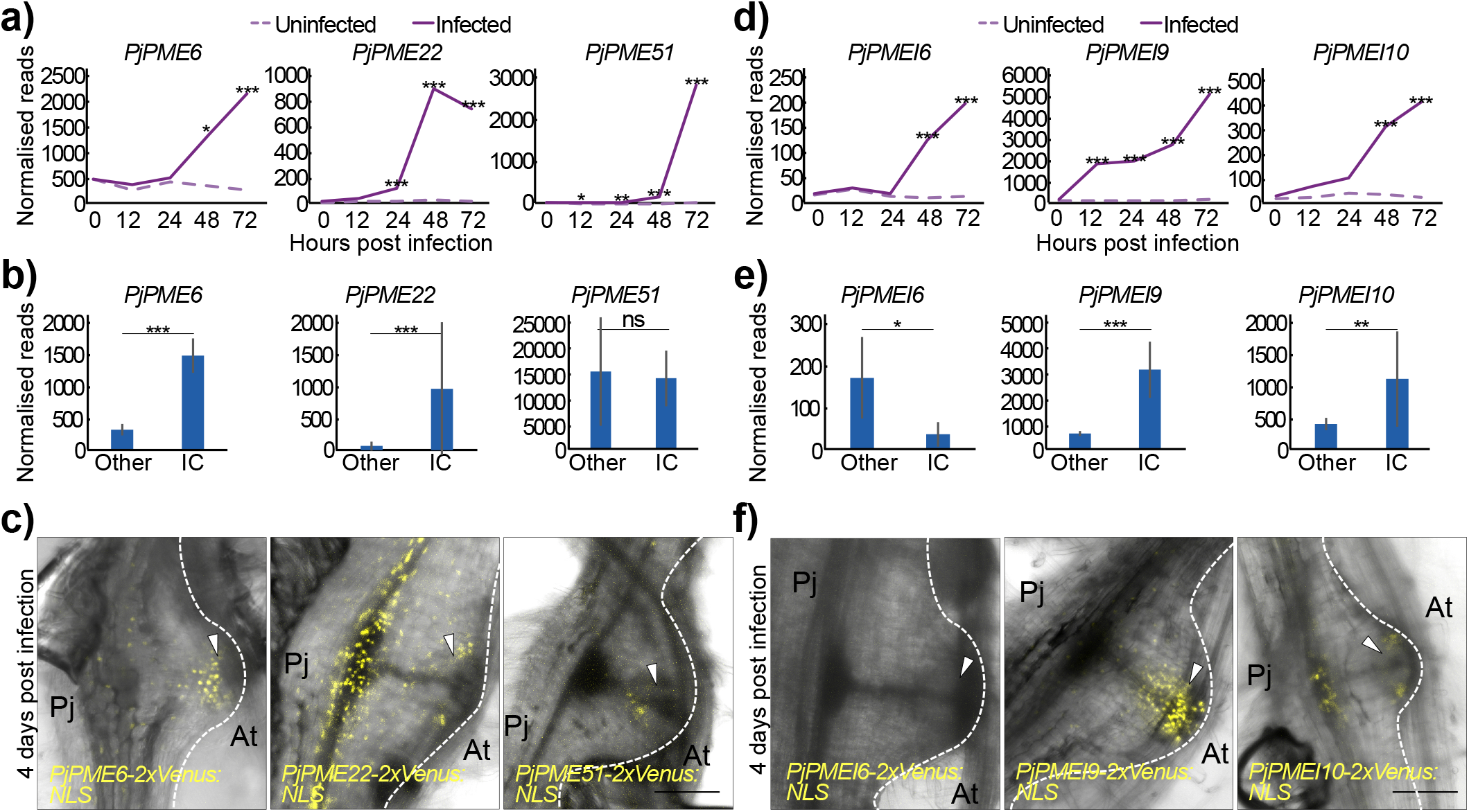
Upregulated *PjPMEs* and *PjPMEIs* are primarily expressed in intrusive cells and cambium-like tissue. a,d) Normalised reads of *PjPME6*, *PjPME22*, *PjPME51, PjPMEI6*, *PjPMEI9* and *PjPMEI10* over five time points during infection for *P. japonicum* infecting and not infecting. Stars indicate a significant difference between infecting and not infecting. b,e) Normalised reads of *PjPME6*, *PjPME22*, *PjPME51, PjPMEI6*, *PjPMEI9* and *PjPMEI10* in intrusive cells and non-IC (other) tissues. Stars indicate a significant difference between IC and other tissues. c,f) Images of infecting transgenic hairy roots expressing *PjPME6*, *PjPME22*, *PjPME51, PjPMEI6*, *PjPMEI9* and *PjPMEI10* nuclear-localised (NLS) transcriptional reporters in fully developed haustoria (4 dpi). Arrowheads denote intrusive cells. Scale bars 100um. * for p<0.05, ** for p<0.01, *** for p<0.001.

### Inhibition of PME activity impairs haustoria induction and development

To determine if PME activity is necessary for haustorium development, we treated infecting *P. japonicum* with epigallocatechin gallate (EGCG), a chemical inhibitor of PME enzymes (Lewis et al. 2008). Treatment reduced the number of haustoria (Fig. 4a) and delayed the formation of the xylem bridge connections to the host (Fig. 4b), although it did not significantly affect plate xylem area and number of xylem bridge connection at 7 dpi (Fig. 4c, Supplementary Fig. 4a,b). EGCG treatment also reduced *PjPME22* and *PjPME51* expression at 72 hpi (Fig. 4d) and reduced *PjPMEI6*, *PjPMEI9* and *PjPMEI10* expression at both 0 and 72 hpi, consistent with chemical inhibition of PMEs affecting both haustoria development and the transcriptional regulation of endogenous *PMEs* and *PMEIs* (Fig. 4d, Supplementary Fig. 4c). We next overexpressed *PjPME6* and *PjPME51* in *P. japonicum* hairy roots (Supplementary Fig. 4d,e) but did not observe defects in haustorium induction or development (Fig. 4e, Supplementary Fig. 4f,g). However, overexpression of *PjPMEI6, PjPMEI9* and *PjPMEI10* with the MAS promoter significantly inhibited haustoria induction (Fig. 4e) but did not affect xylem connections (Supplementary Fig. 4f,g). Finally, we tested whether modifying pectin status in the host could affect infection by using the *PMEI5*-overexpressing *A. thaliana* line *AtPMEI5OE*, which is characterised by highly methylesterified pectin (Wolf et al. 2012; Jonsson et al. 2021). Haustoria induction was not affected in the mutant compared to wild-type Col-0 (Fig. 4f), however, xylem bridge formation was delayed during infection of *AtPMEI5OE* (Fig. 4g). Taken together, these results suggest that parasitic PME activity is important for efficient induction and development of haustoria.

**Figure 4:**
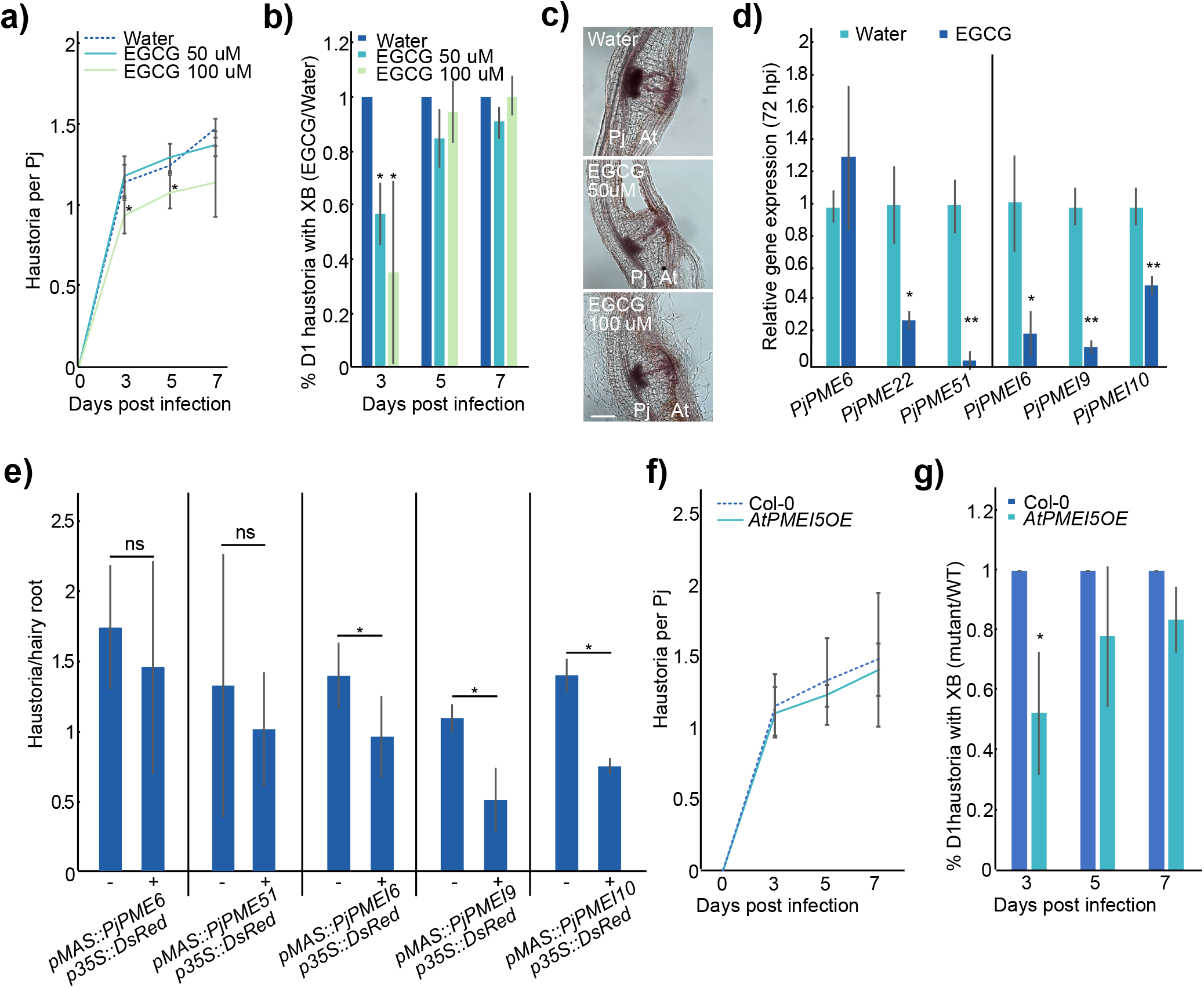
Inhibition of PME activity impairs haustorium induction and development. a) Number of haustoria per *P. japonicum* plant at four time points during treatment with 50 or 100 uM EGCG or water control. Asterisks indicate significance compared to control (Student’s t-test) b) Ratio of the percentage of day-1 (D1) haustoria with a xylem bridge formed during treatment with 50 or 100 uM EGCG over water at three time points. Asterisks indicate significance compared to control (Student’s t-test) c) Images of 7 dpi haustoria formed on water, EGCG 50 uM and EGCG 100 uM. d) Relative gene expression of selected *PjPMEs* and *PjPMEIs* at 72 hpi in *P. japonicum* haustoria treated with 100 uM EGCG, normalised to water. Asterisks indicate significance compared to control (Student’s t-test) e) Numbers of haustoria per hairy root forming on hairy roots transformed with *PjPME* and *PjPMEI* overexpression constructs. Non-transgenic hairy roots are marked as “−” and transgenic roots are marked as “+”. Asterisks indicate significance compared to control (Fisher exact test) f) Number of haustoria per *P. japonicum* plant at four time points during infection of *AtPMEI5OE* mutant or Col-0 as control. g) Ratio of the percentage of D1 haustoria with a xylem bridge formed during infection of *AtPMEI5OE* over Col-0 at three time points. Asterisks indicate significance compared to control (Student’s t-test). * for p<0.05, ** for p<0.01.

### Brassinosteroid treatment reduces *PjPME* and *PjPMEI* expression and delays haustorium development

Brassinosteroid (BR) signalling mediates cell wall biosynthesis and remodelling, and has been implicated in feedback mechanisms with PME and PMEI activity and EGCG treatments (Wolf et al. 2012). To test the effect of BRs, we applied epibrassinolide (epiBL) during *P. japonicum* infection and found it reduced the number of haustoria per *P. japonicum* and inhibited the formation of xylem bridges (Fig. 5a,b, Supplementary Fig. 5a), similar to EGCG treatment (Fig. 4a,b). We also tested the expression of *PjPME*s and *PjPMEI*s in haustoria following epiBL treatment. *PjPME51*, *PjPMEI6*, *PjPMEI9* and *PjPMEI10* were downregulated in haustoria treated with epiBL at 72 hpi but not at 0 hpi (Fig. 5c, Supplementary Fig. 5b), suggesting transcriptional control of pectin methylesterification by BR signalling during haustorium development. To determine if BR treatment could modify pectin methylesterification levels, we performed antibody staining using LM19 and LM20 on cross sections of 0 hpi and 72 hpi haustoria untreated or treated with epiBL (Fig. 5d, Supplementary Fig. 5c). The haustoria treated with epiBL had lower levels of both unmethylesterified pectin (LM19) and highly methylesterified pectin (LM20) compared to the control (Fig. 5e), suggesting overall pectin levels were reduced following epiBL treatment. Finally, we infected *A. thaliana* mutants with modified BR signalling, *bes1-2* and *bes1-D*, to test the role of host BR signalling during infection. *P. japonicum* could efficiently infect both mutants and establish xylem connections (Supplementary Fig. 5d,e), suggesting host BR signalling is not crucial for haustoria development and instead BR signalling might be important for parasite cell wall modifications during infection.

**Figure 5:**
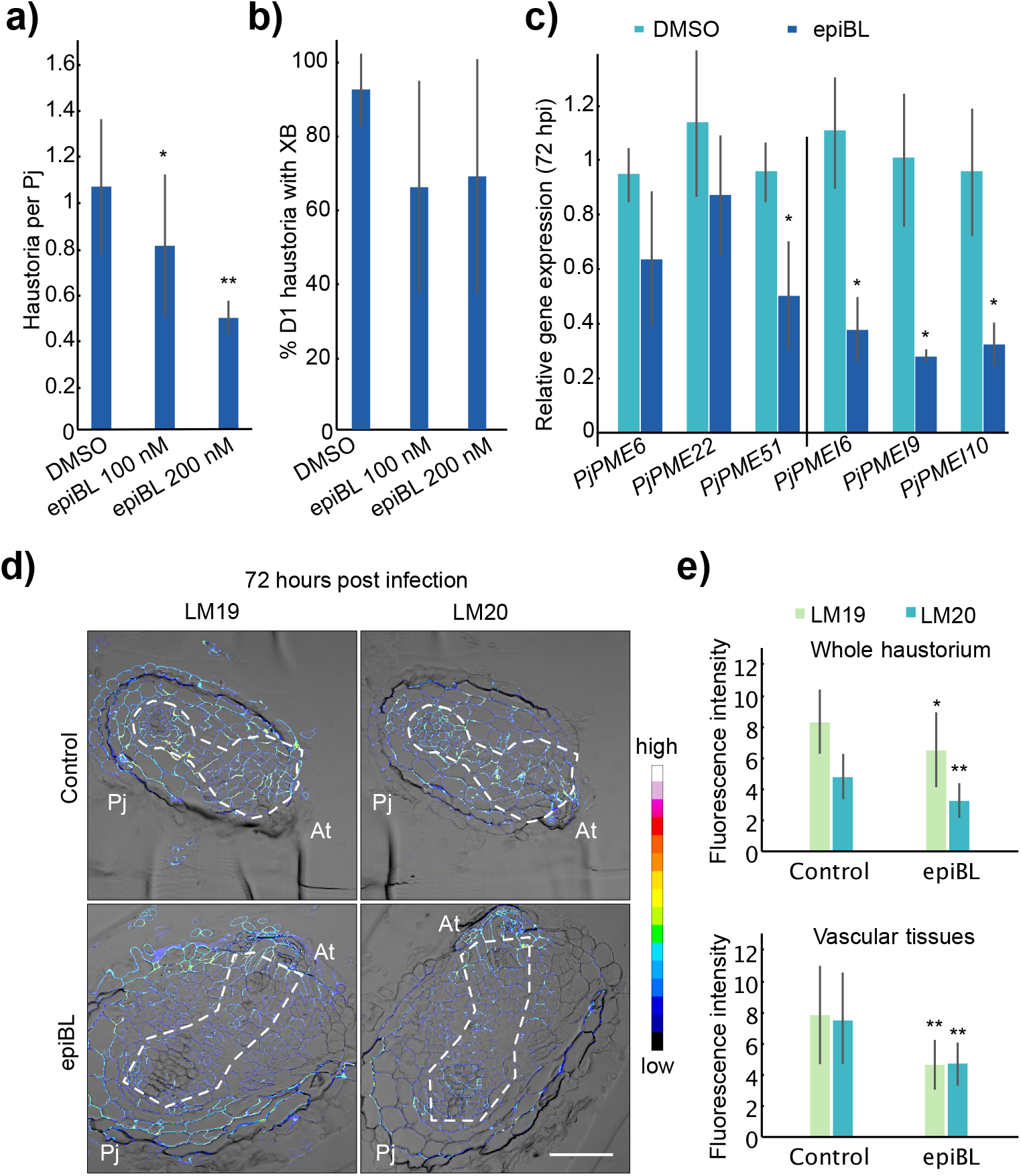
BR treatment reduces *PjPME* and *PjPMEI* expression and inhibits haustorium development. a) Number of haustoria per *P. japonicum* plant at 7 dpi during treatment with 100 or 200 nM epiBL or DMSO control. b) Percentage of day-1 (D1) haustoria with a xylem bridge formed during treatment with 100 or 200 nM epiBL or DMSO at 7 dpi. c) Relative gene expression of selected *PjPMEs* and *PjPMEIs* at 72 hours post infection in *P. japonicum* haustoria treated with 100 nM epiBL, normalised to DMSO. d) Fluorescence images of antibody staining using LM19 (unmethylated homogalacturonan) and LM20 (highly methylated homogalacturonan) on cross sections of 72 hpi haustoria developed on DMSO control or 100 nM epiBL. Scale bars 100 um. Vascular tissues are circled in white dashed lines. e) Fluorescence quantification for whole *P. japonicum* sections and vascular tissues. Asterisks indicate significance compared to control (Student’s t-test): * for p<0.05, ** for p<0.01.

### *PjPMEs* and *PjPMEIs* expression correlates with xylem bridge development

Since some *PjPMEs* and *PjPMEIs* are expressed in the cambium and xylem-like tissues during haustorium development (Fig. 3c-f) and inhibiting PME activity delays xylem formation (Fig. 4b), we investigated the role of PjPMEs and PjPMEIs during xylem-bridge formation. Looking at the expression pattern of cambium and xylem marker genes (Kokla et al. 2022), we found the cambium marker *PjWOX4* (Wakatake et al. 2018) co-expressed with *PjPME22*, *PjPME51* and *PjPMEI9* (Fig. 6a). The procambium marker *PjHB8* co-expressed with *PjPME6*. The xylem-markers *PjCESA7* (Wakatake et al. 2018), *PjVND7* (identified through BLAST using AtVND7 as a query) and *PjXCP2* (Kokla et al. 2022) co-expressed with *PjPMEI6* and *PjPMEI10* (Fig. 6a, Supplementary Fig. 6a). To investigate if any of these *PjPMEs* and *PjPMEIs* are expressed in response to xylem-bridge development, we chemically inhibited xylem bridge formation by treatment with the auxin transport inhibitor N-1-naphthylphthalamidic acid (NPA). NPA treatment did not affect haustoria numbers, yet completely inhibited xylem bridge connection (Wakatake et al. 2020, Fig. 6b-d). Treatment with the synthetic auxin 1-naphthaleneacetic acid (NAA) did not affect haustoria numbers or xylem bridge formation (Supplementary Fig. 6b,c), but increased expression of *PjPMEI9* and *PjPMEI10* in non-infecting *P. japonicum* roots, and decreased *PjPME22* and *PjPMEI9* expression in 72 hpi haustoria (Supplementary Fig. 6d). We also found that the commonly used dye Coomassie Brilliant Blue, an inhibitor of xyloglucan endotransglucosylase/hydrolase (XTH) activity (Olsen and Krause 2017), increased haustoria numbers yet reduced xylem bridge formation by approximately 60% (Fig. 6b-d). LM19 and LM20 antibody staining of 72hpi sections showed both de-methylated pectin and highly methylated pectin were reduced following NPA and Coomassie treatments (Fig. 6e,f, Supplementary Fig. 6e). Expression levels of *PjPME51* and *PjPMEI9* were also significantly decreased by both treatments at 72 hpi (Fig. 6g) but not at 0 hpi (Supplementary Fig. 6f). These data indicated that decrease expression of *PjPME51* and *PjPMEI9* was specific to xylem bridge inhibition and suggested a role for these genes in xylem bridge formation. We then infected hairy roots expressing *PjPMEI9-2xVenus:NLS*, which showed fluorescence in cambium-like tissues (Fig. 3f, Supplementary Video1). At 4 days post infection we observed a marked decrease in fluorescence when hairy roots expressing *PjPMEI9-2xVenus:NLS* were treated with NPA compared to the control, consistent with the qPCR data (Fig. 6h) and demonstrating that xylem bridge formation is important for *PMEI9* expression.

**Figure 6:**
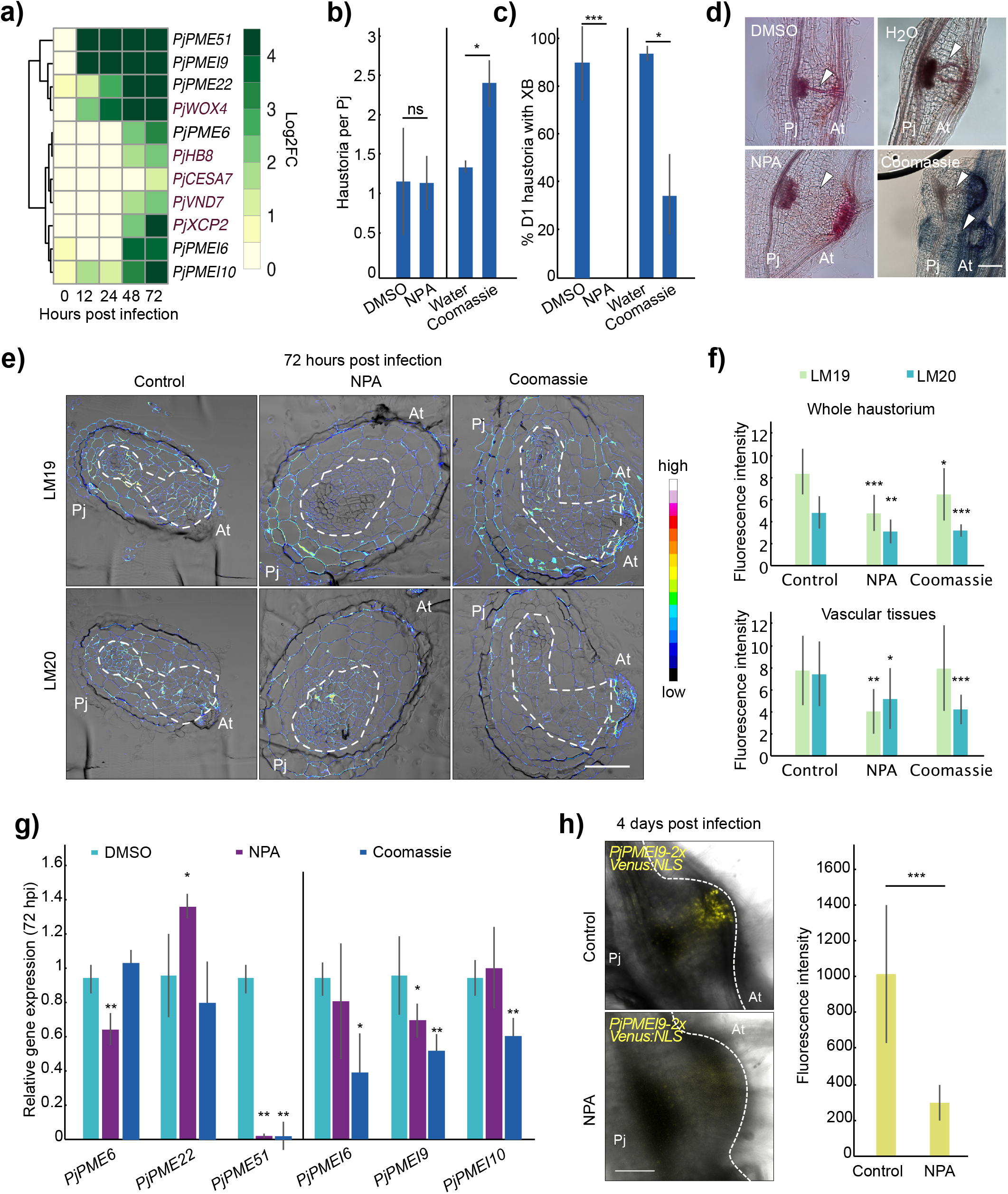
*PjPMEs* and *PjPMEIs* expression correlates with xylem bridge development. a) Expression heatmap of *P. japonicum* cambium and xylem marker genes (red) and selected *PjPMEs* and *PjPMEIs*: log2 fold change between *P. japonicum* infecting and not infecting over five time points during infection b) Number of haustoria per *P. japonicum* plant at 7 dpi during treatment with DMSO, 5 uM NPA, water or 0.05 mM Coomassie. c) Percentage of day-1 (D1) haustoria with a xylem bridge formed during treatment with DMSO, 5 uM NPA, water or 0.05 mM Coomassie. d) Images of 7 dpi haustoria formed on DMSO, NPA 5 uM, water or 0.05 mM Coomassie. White arrowheads denote the location of XB development e) Fluorescence images of antibody staining using LM19 (unmethylated homogalacturonan) and LM20 (highly methylated homogalacturonan) on cross sections of 72 hpi haustoria developed on DMSO control, 5 uM NPA or 0.05 mM Coomassie. Vascular tissues are circled in white dashed lines. f) Fluorescence quantification for whole *P. japonicum* haustoria and vascular tissues. g) Relative gene expression of select *PjPMEs* and *PjPMEIs* at 72 hpi in *P. japonicum* haustoria treated with 5 uM NPA or 0.05 mM Coomassie normalised to DMSO. h) Images of haustoria developed on hairy roots expressing the *PjPMEI9* nuclear-localised (NLS) transcriptional reporter at 4 dpi on DMSO or 5 uM NPA, and quantification of fluorescence intensity. Scale bars 100 um. Asterisks indicate significance compared to control (Student’s t-test): * for p<0.05, ** for p<0.01, *** for p<0.001.

## Discussion

### Haustoria induction and development in *P. japonicum* require PME activity from the parasite

Here, we investigated the role of PME-mediated pectin modifications during haustorium development in *P. japonicum* and identified multiple PMEs and their inhibitors that were upregulated during both *Arabidopsis* and rice infections. We found that PME activity, and likely the demethylesterification of pectins, was needed for efficient haustorium induction and host penetration since inhibition of PME activity through EGCG treatment or PMEI overexpression in the parasite reduced haustoria numbers (Fig. 4a,e). Recently, it was found that pectin methylesterification levels are also crucial for lateral root initiation (Wachsman et al. 2020). Our findings are therefore consistent with aspects of lateral root and haustoria emergence being conserved, as previously suggested in *Cuscuta* and *Thesium* (Ichihashi et al. 2018; Jhu et al. 2021). We also observed a decrease in pectin methylesterification at 48 and 72 hpi, however, this did not occur in DMBQ-induced pre-haustoria (Fig. 2a,b, Supplementary Fig. 2a,b) and it is likely that PME activity was specifically linked to host penetration by *P. japonicum*. There was a strong increase in *PMEs* and *PMEIs* expression by 48-72 hpi (Fig. 1c,d, 3a,d) and this increase of both enzyme and inhibitor could be explained by different spatial expression, leading to the different degrees of pectin methylesterification (DMs) in different tissues (Fig. 2c, Supplementary Fig. 2c). Asymmetric pectin DM is required in several plant developmental processes, including apical hook formation during seedling emergence and leaf patterning (Jonsson et al. 2021; Peng et al. 2022). Furthermore, the difference in pectin DM between lateral and primary roots might allow the emergence of the lateral root while preventing its own digestion (Laskowski et al. 2006). We propose that in haustoria the high PME activity (low DM) in intrusive cells drives host cell wall loosening and penetration, while the high PMEI activity (high DM) is required in inner haustorial tissues to maintain organ integrity and push towards the host. In addition to PMEs, the subtilase enzyme family are also expressed in the intrusive cells of developing haustoria and are important for xylem bridge development in *P. japonicum* (Ogawa et al. 2021). Subtilases can control PME activity by cleaving PME proteins to the final active enzyme form (Senechal et al. 2014). Several haustoria-specific *PjPMEs* we investigated were also expressed in intrusive cells (Fig. 3b,c) and we speculate that subtilases might process some or all of these PjPMEs to allow host penetration and efficient vascular connection to the host. More studies will however be needed to confirm this hypothesis. In the *Arabidopsis* host, our data suggest the role of pectin modification during infection is less important. Only *AT1G23200* (*PME*) and *AT2G01610* (*PMEI*) showed clear increases in expression during infection (Supplementary Fig. 1c,d), suggesting these genes might either be involved in the defence response to the parasite or activated by *P. japonicum* to facilitate parasitism. Furthermore, infection of the *Arabidopsis* overexpressor *AtPMEI5OE* (Wolf et al. 2012), which has high DM (Wolf et al. 2012; Jonsson et al. 2021) did not affect the total number of haustoria and only delayed xylem bridge connection (Fig. 4f,g). Taken together, our results point to PME-mediated pectin modifications being most relevant for the parasite during haustorium initiation and host invasion.

### Brassinosteroid and auxin and signalling pathways interact with pectin modifications during haustorium development

We found that exogenous epiBL treatment of *P. japonicum* partially inhibited haustoria induction (Fig. 5a) and reduced the expression of several *PjPME* and *PjPMEI* genes during infection (Fig. 5c). Brassinosteroid signalling is implicated in controlling cell-wall modifications. There, modification of pectin DM activates brassinosteroid signalling which can then modulate PME activity, resulting in a feedback loop to control cell-wall loosening (Wolf et al. 2012; Wolf et al. 2014). Our data suggest that some *PjPMEs* and *PjPMEIs* are responsive to BR signalling (Fig. 5c), and epiBL application reduced both methylesterified and de-methylesterified pectins in haustoria (Fig. 5d,e). BR signalling might therefore regulate PME activity also during haustorium development.

High auxin also promotes pectin loosening (Braybrook and Peaucelle 2013). Our NAA treatments induced *PjPMEI9* and *PjPMEI10* in non-infecting *P. japonicum* roots, suggesting these genes are activated by auxin signalling (Supplementary Fig. 6d). Furthermore, blocking auxin transport through NPA treatment reduced *PjPMEI9* expression during haustorium development (Fig. 6g,h). These data indicate that auxin transported to the haustorium site during development likely induces *PjPMEI9* expression. In *P. japonicum*, haustoria initiation requires local auxin biosynthesis (Ishida et al. 2016) and auxin transport drives xylem bridge development (Wakatake et al. 2020). Previous studies have also shown that high auxin levels trigger pectin de-methylesterification during differential *Arabidopsis* seedling growth, and increase PME expression during lateral root emergence (Laskowski et al. 2006; Jonsson et al. 2021). We hypothesise that auxin signalling plays a crucial role during haustorium formation by regulating pectin loosening and PME activity to allow haustorium emergence, host penetration and tissue differentiation.

### Pectin modifications affect xylem bridge differentiation

Xylem bridge-inhibiting treatments reduced pectin levels and caused differential expression of *PjPMEs* and *PjPMEIs* during haustoria development (Fig. 6e-h). In addition, blocking PME activity reduced xylem bridge formation (Fig. 4b). Taken together, these results suggest a role for PME activity during xylem bridge differentiation. In *Medicago sativa*, xylem cell walls contain about 4% pectin, compared to the 25% of other tissues (Grabber et al. 2002), suggesting pectin might be degraded during xylem differentiation. In *Arabidopsis*, five *PMEs* are expressed in xylem tissues (Pelloux et al. 2007), and the demethylesterification of pectin might be important for lignification (Lairez et al. 2005; Pelloux et al. 2007). PME activity might therefore be required in the first stage of xylem bridge differentiation to allow pectin degradation, followed by lignification. In *P. japonicum*, we observed increased PME activity at 48 and 72 hpi, when the xylem plate and xylem bridges start developing (Fig. 2a,b). Our data also suggest *PjPME51* and *PjPMEI9* might be involved in xylem bridge development, as they are expressed in these tissues (Fig. 3c,f) and their expression level is reduced by inhibiting xylem bridge formation (Fig. 6g,h). *PjPME51* might contribute to the demethylesterification of the developing xylem strands, allowing for pectin degradation and secondary cell wall development. *PjPMEI9* might instead be maintaining a high level of methylesterification in the surrounding tissues (Fig. 2c,d), improving tissue stability and limiting the action of PjPME51 and possibly other PMEs only to the tissues where it is required. Thus, our data suggest a strong link between pectin modifications and xylem development but further investigations will be needed to test this relationship in non haustorial tissues and also the role of *PjPME51* and *PjPMEI9*.

## Acknowledgements

We thank Sebastian Wolf for providing the *AtPMEI5OE* seeds, Thomas Spallek for providing the GoldenGate plasmid containing the 3xVenus-NLS sequence and Judith Lundberg-Felten for kindly donating the LM19 and LM20 antibodies. M.L., M.F., and C.W.M. were supported by an ERC starting grant (GRASP-805094). A.K. and C.W.M were supported by a Wallenberg Academy Fellowship (2016-0274). M.F. was supported by a Formas Grant (2018-00533).

## Author Contributions

ML and CWM conceived the experiments. ML, AK and MF performed the experiments. ML and CWM wrote the manuscript. All authors revised the final manuscript.

## Supplementary material

Supplementary Fig. 1: *Arabidopsis PMEs* and *PMEIs* are differentially expressed during haustorium development

Supplementary Fig. 2: PME activity is increased during haustorium development

Supplementary Fig. 3: *PjPMEs* and *PjPMEIs* are specific to haustoria

Supplementary Fig. 4: *PjPME* and *PjPMEI* overexpression does not affect xylem connection to the host

Supplementary Fig. 5: Arabidopsis BR signalling mutants do not affect parasitism efficiency

Supplementary Fig. 6: NAA treatment does not affect parasitism efficiency

Supplementary Table 1: List of identified PjPMEs and PjPMEIs

Supplementary Table 2: *P. japonicum* gene accessions and IDs Supplementary

Table 3: Primers used in the study

SupplemetaryVideo1: Z-stack of a haustorium expressing *PjPMEI9-2xVenus∷NLS*

## Declaration of interests

The authors declare no competing interests

## Notes

### Competing Interest Statement

The authors have declared no competing interest.

