## Supplemental Figures 1-6 for "PME-mediated pectin modifications promote haustoria initiation and xylem bridge development in the parasitic plant *Phtheirospermum japonicum*"

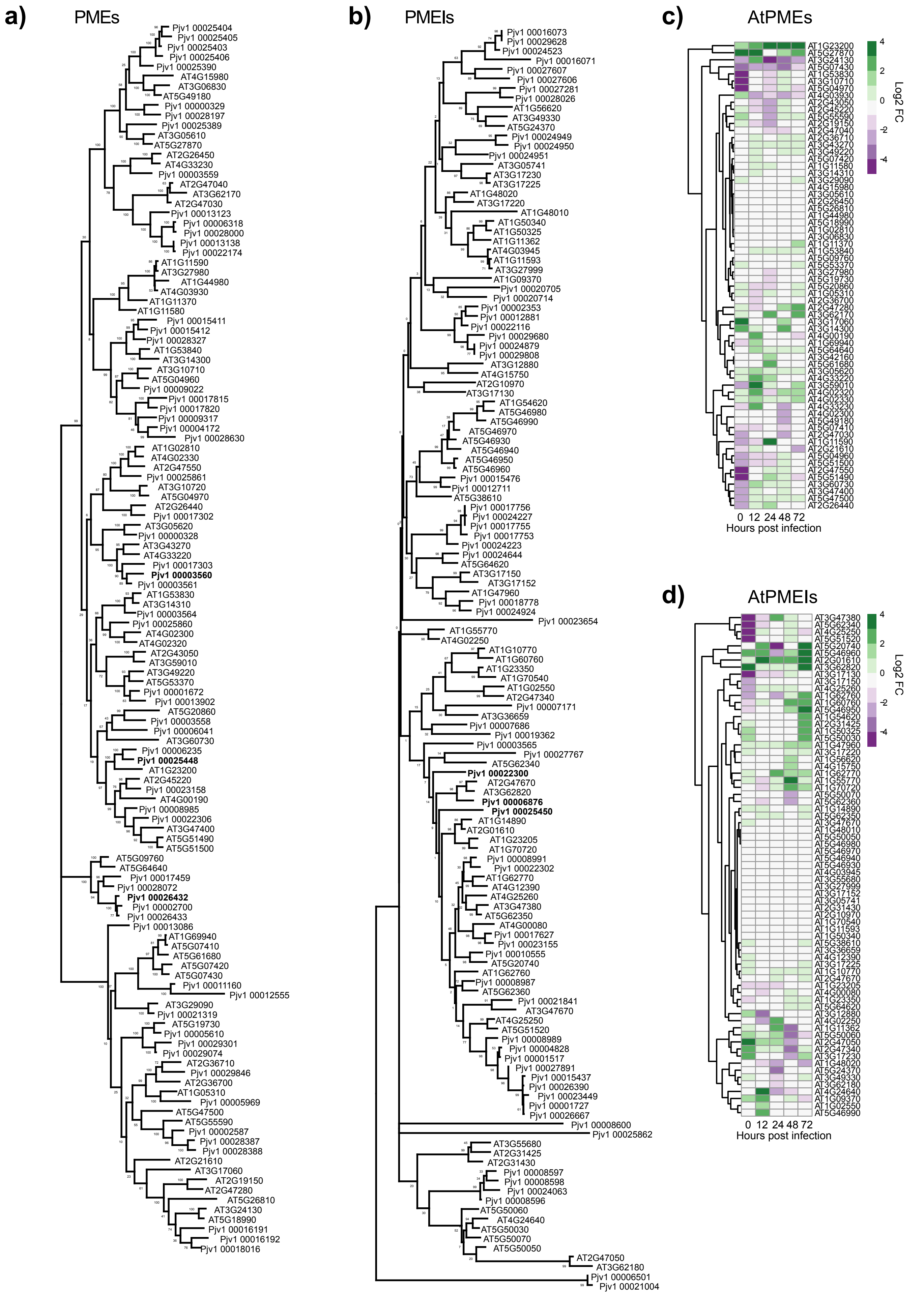


**S1: *Arabidopsis PMEs* and *PMEIs* are differentially expressed during haustorium development**
a) Maximum-Likelihood phylogenetic tree of PjPMEs and AtPMEs. b) Maximum-Likelihood phylogenetic tree of PjPMEIs and AtPMEIs c) Heatmap of the expression of *A. thaliana PMEs*: log2 fold change between infected and not infected *Arabidopsis* over five time points during infection d) Heatmap of the expression of *A. thaliana PMEIs*: log2 fold change between infected and not infected *Arabidopsis* over five time points during infection

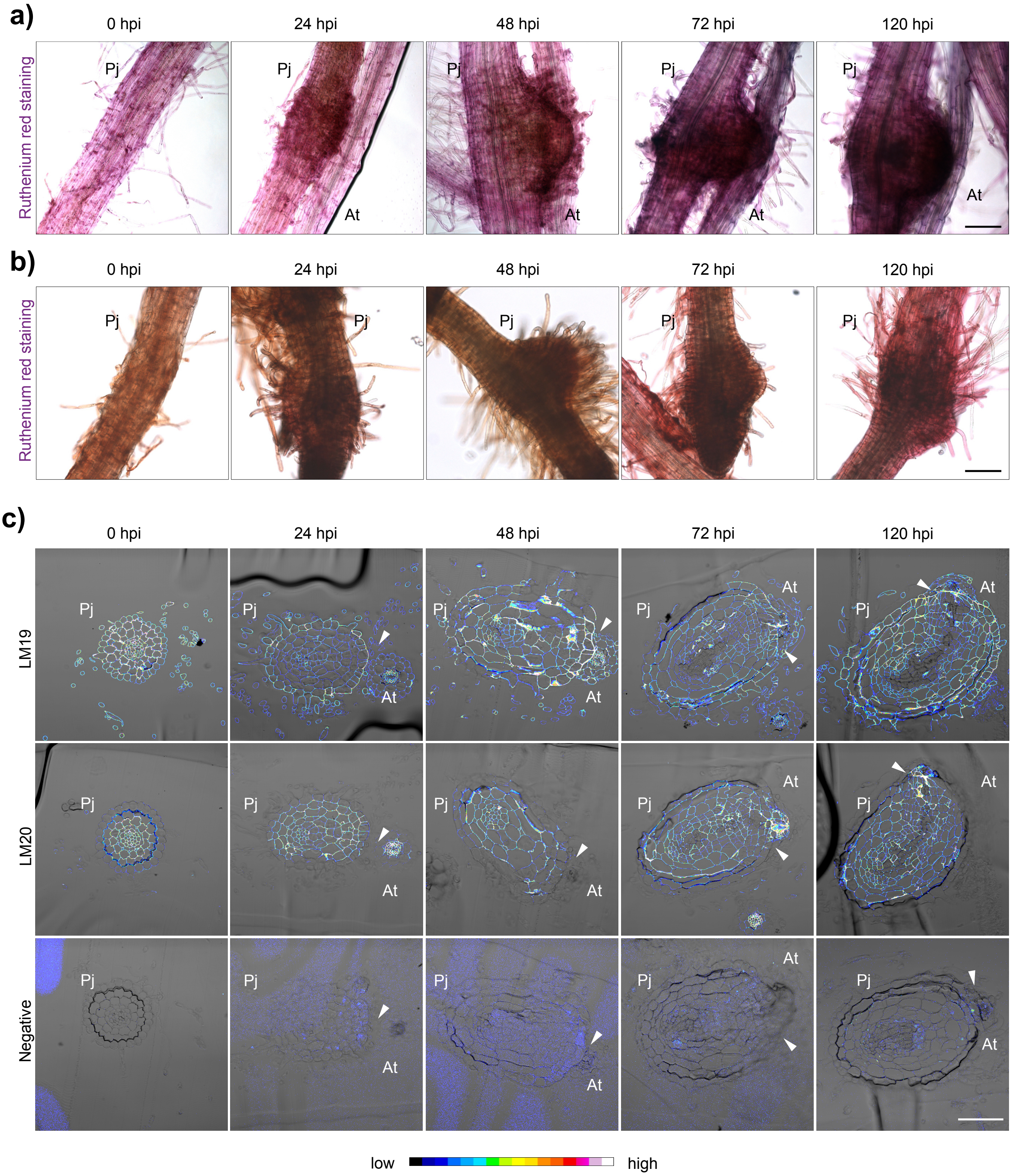


**S2: PME activity is increased during haustorium development**
a) Ruthenium red staining of the developing haustoria at 0, 24, 48, 72 and 120 hours post infection (hpi). 24, 48 and 120 hpi images are repeated from Fig. 2a b) Ruthenium red staining of the developing pre-haustorium at 0, 24, 48, 72 and 120 hours post exposure to DMBQ. c) Fluorescence images of antibody staining using LM19 (unmethylated homogalacturonan), LM20 (highly methylated homogalacturonan) and negative control (PBS) on haustoria cross section at 0, 24, 48, 72 and 120 hpi. 24, 48 and 120 hpi images for LM19 and LM20 are repeated from Fig. 2b. Scale bars 100 um; Pj = *Phtheirospermum japonicum*, At = *Arabidopsis thaliana*; arrowheads point at the parasite-host interface.

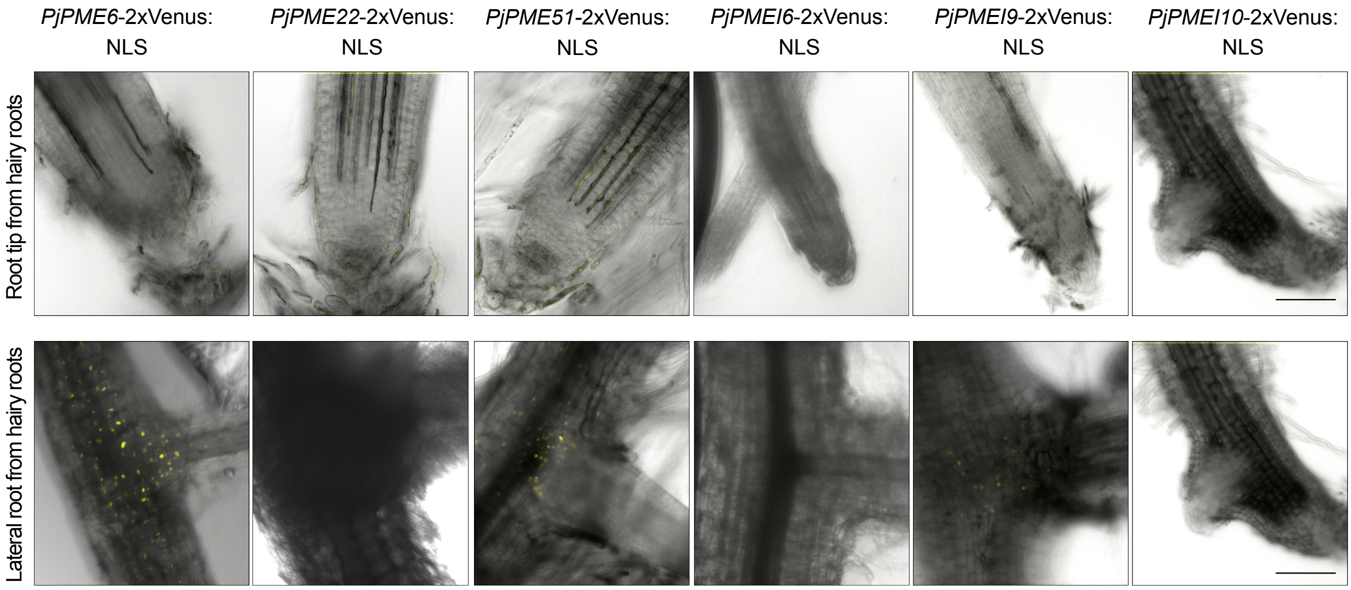


**S3: *PjPMEs* and *PjPMEIs* are specific to haustoria**
Images of transgenic hairy roots expressing *PjPME6*, *PjPME22, PjPME51, PjPMEI6*, *PjPMEI9* and *PjPMEI10* nuclear-localised transcriptional reporters: root tips and lateral roots.
Scale bars 100 um.

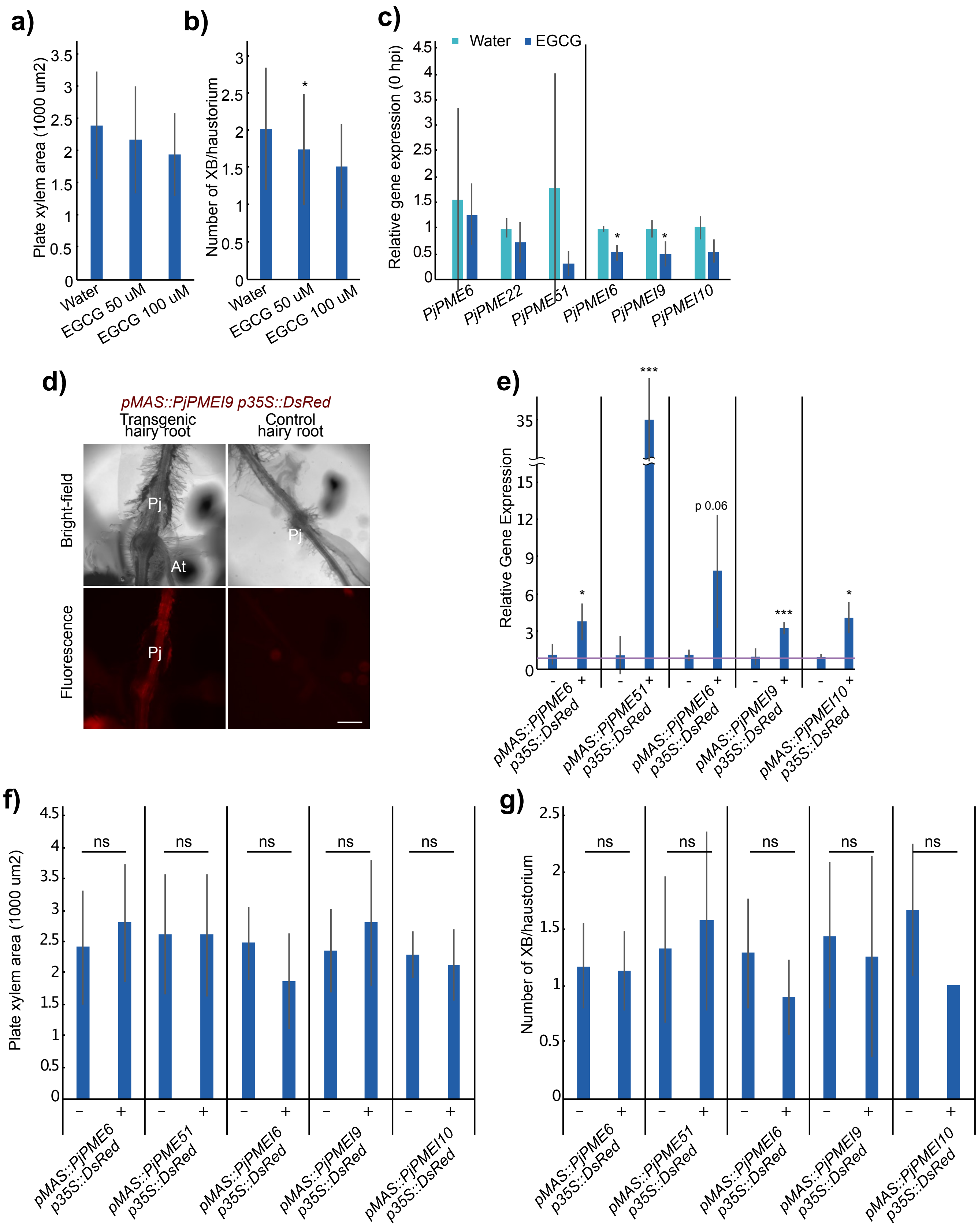


**S4: *PjPME* and *PjPMEI* overexpression does not affect xylem connection to the host**
a) Area of plate xylem in 7 dpi haustoria treated with 50 or 100 uM EGCG or water as control. b) Number of xylem bridges per haustorium in 7 dpi haustoria treated with 50 or 100 uM EGCG or water as control. c) Relative gene expression of selected *PjPMEs* and *PjPMEIs* at 0 hpi in *P. japonicum* haustoria treated with 100 uM EGCG, normalised to water. d) Representative images of hairy roots transformed with gene overexpression constructs. Control hairy roots show no fluorescence, while transformed hairy roots have red fluorescence. Scale bar 400 um. e) Relative gene expression of the *PjPME* or *PjPMEI* of interest in hairy roots transformed with the indicated construct (+) and control roots (-). f) Area of plate xylem in 7 dpi haustoria formed on hairy roots transformed with the indicated construct (+) and control roots (-). g) Number of xylem bridges per haustorium in 7 dpi haustoria formed on hairy roots transformed with the indicated construct (+) and control roots (-). Asterisks indicate significance compared to control (Student's t-test)* for $p<0.05$, ** for $p<0.01$, *** for $p<0.001$.

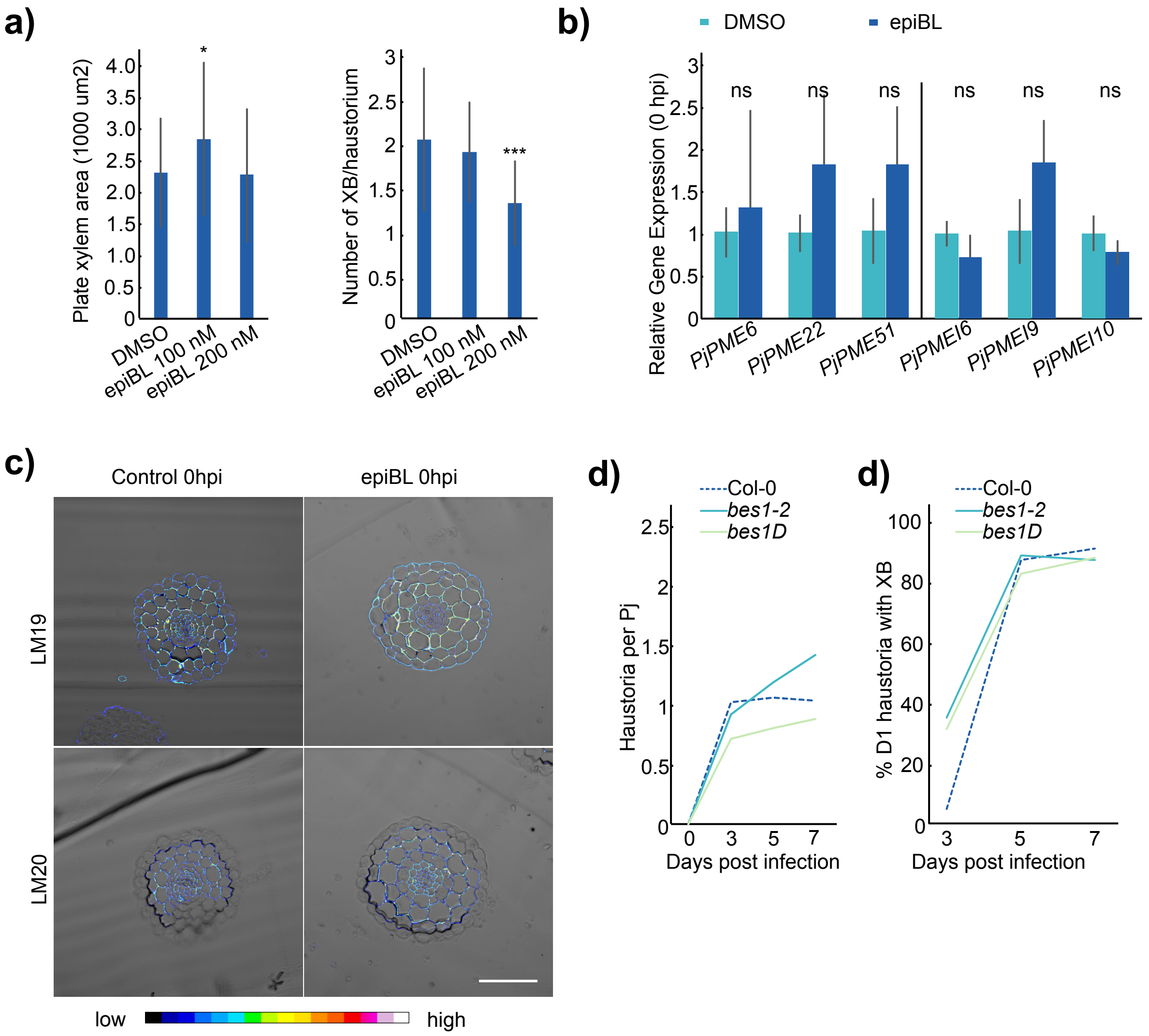


**S5: Arabidopsis BR signalling mutants do not affect parasitism efficiency**
a) Area of plate xylem in 7 dpi haustoria treated with 100 nM and 200 nM epiBL or DMSO as control and number of xylem bridges per haustorium in 7 dpi haustoria treated with 100 and 200 nM epiBL or DMSO at 7 dpi. b) Relative gene expression of selected *PjPMEs* and *PjPMEIs* at 0 hpi in *P. japonicum* haustoria treated with 100 nM epiBL, normalised to DMSO. c) Fluorescence images of antibody staining using LM19 (unmethylated homogalacturonan) and LM20 (highly methylated homogalacturonan) on cross sections of haustoria developed on DMSO (Control) or 100 nM epiBL at 0 hpi. Scale bar 100 um. d) Number of haustoria per *P. japonicum* plant at four time points during infection of *bes1-2* and *bes1-D* mutants or Col-0 as control. e) Percentage of Day-1 (D1) haustoria with a xylem bridge formed during infection of *bes1-2* and *bes1-D* mutants or Col-0 at three time points; n=2. Asterisks indicate significance compared to control (Student's t-test): * for p<0.05, ** for p<0.01, *** for p<0.001.

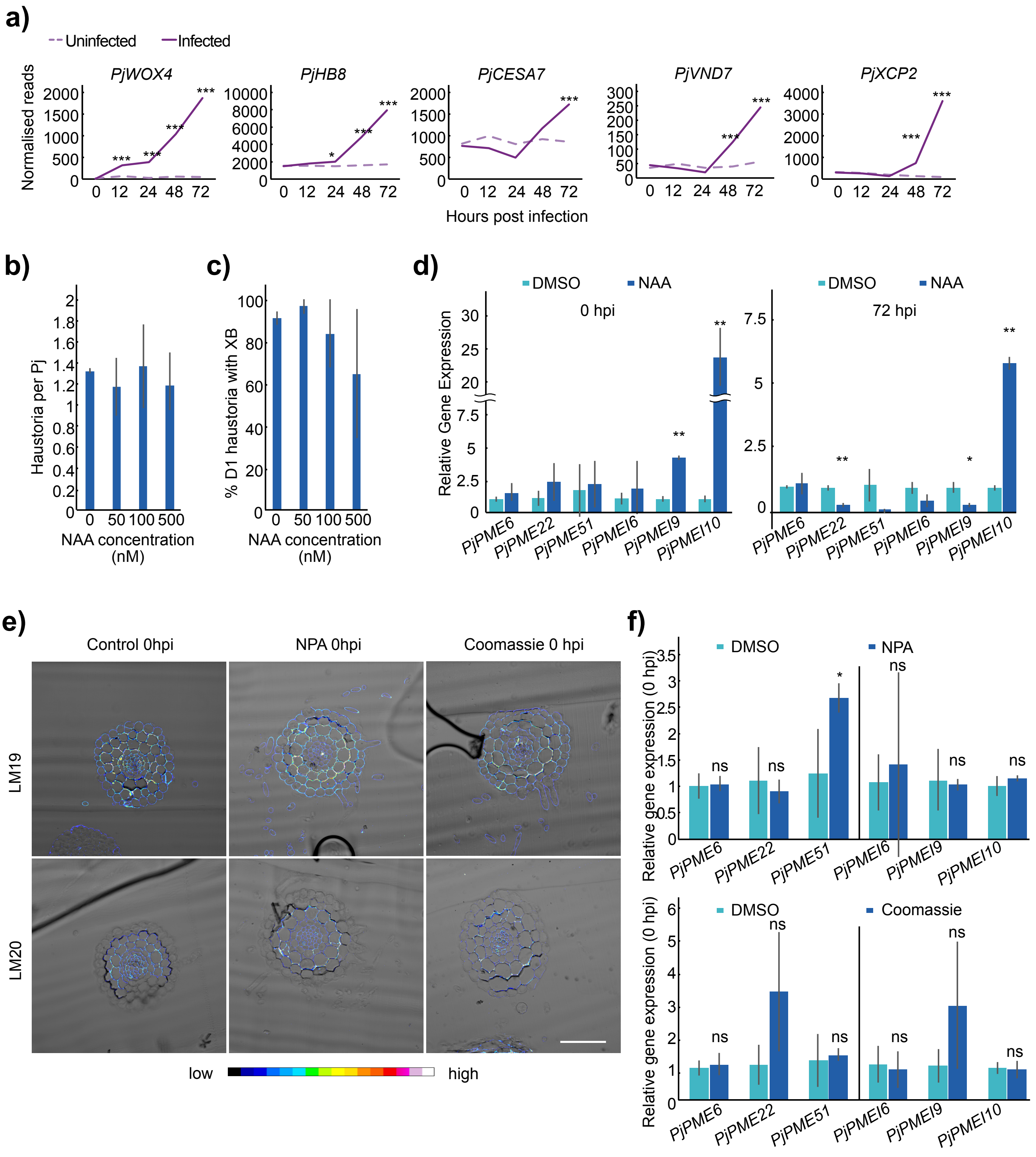


**S6: NAA treatment does not affect parasitism efficiency**

a) Normalised reads of *PjWOX4*, *PjHB8*, *PjCESA7*, *PjVND7* and *PjXCP2* over five time points during infection for *P. japonicum* infecting and not infecting. Stars indicate significant difference between infecting and not infecting. b) Number of haustoria per *P. japonicum* plant at 7 dpi during treatment with 0, 50, 100 or 500 nM NAA c) Percentage of Day-1 (D1) haustoria with a xylem bridge formed during treatment with 0, 50, 100 or 500 nM NAA. d) Relative gene expression of selected *PjPMEs* and *PjPMEIs* at 0 and 72 hours post infection in *P. japonicum* haustoria treated with 1 uM NAA, normalised to DMSO. e) Fluorescence images of antibody staining using LM19 (unmethylated homogalacturonan) and LM20 (highly methylated homogalacturonan) on cross sections of 0 hpi haustoria developed on DMSO (Control), 5 uM NPA or 0.05 mM Coomassie BB. Scale bar 100 um. f) Relative gene expression of selected *PjPMEs* and *PjPMEIs* at 0 hpi in *P. japonicum* haustoria treated with 5 uM NPA, normalised to DMSO, or 0.05 mM Coomassie BB, normalised to DMSO.
